# Rapid dissolution of dioecy by experimental evolution

**DOI:** 10.1101/712414

**Authors:** G. G. Cossard, J. F. Gerchen, X. Li, Y. Cuenot, J. R. Pannell

## Abstract

Evolutionary transitions from hermaphroditism to dioecy have been frequent in flowering plants, but recent analysis indicates that reversions from dioecy to hermaphroditism have also been common. Here, we use experimental evolution to expose a mechanism for such reversions. We removed males from dioecious populations of the wind-pollinated plant Mercurialis annua and allowed natural selection to act on the remaining females that varied in their propensity for the occasional production of male flowers; such ‘leaky’ sex expression is common in both males and females of dioecious plants. Over only four generations, females evolved a 23-fold increase in average male-flower production. The phenotypic masculinization of females was also sufficient to render them effective at siring progeny in the presence of males. Our study illustrates the rapid dissolution of dioecy and the evolution of functional hermaphroditism under conditions that may frequently occur during periods of low population density, repeated colonization, or range expansion. It thereby experimentally validates a mechanism for a major transition in plant sexual systems..

---

Dioecy (the occurrence of separate sexes) is found in approximately half of flowering plant families, but dioecious species account for only about 6% of flowering plant diversity (*1, 2*). This scattered phylogenetic distribution of separate sexes was long thought to imply that dioecy is an evolutionary dead end, leading to reduced diversification or increased extinction (*3-5*). Recent comparative analysis, however, has concluded that the evolution of separate sexes promotes greater diversification, with no increase in the risk of extinction (*6-8*), casting doubt in the ‘dead-end’ hypothesis. Rather, dioecy might be rare and phylogenetically scattered because of reversions to hermaphroditism (*8, 9*). This ‘reversion’ hypothesis is consistent with the inferred dioecious ancestry for several large predominantly dioecious plant families (*8, 9*). However, reversion to hermaphroditism from separate sexes via the breakdown of dioecy has never been experimentally demonstrated.

The breakdown of dioecy involves either the evolution of functionally bisexual flowers (with both carpels and stamens), or the evolution of bisexual individuals that produce separate male and female flowers on the same plant (monoecy) (*10*). The strong phylogenetic association of dioecy with monoecy (*1*) might thus be due to frequent transitions between monoecy and dioecy in both directions. Transitions from monoecy to dioecy are thought to be due to selection for sexual specialization and/or inbreeding avoidance (*11, 12*), but how might the reverse transition from dioecy to monoecy or hermaphroditism occur? Here, we show that strong mate limitation in experimental populations of the dioecious plant *Mercurialis annua* brings about the rapid breakdown of dioecy via the evolution of masculinized females with dramatically enhanced male-flower production. *Mercurialis annua* is an annual wind-pollinated ruderal weed with XY sex determination (*13, 14*) and strong dimorphism in inflorescence architecture (Fig. 1a and b).

**Fig. 1.**
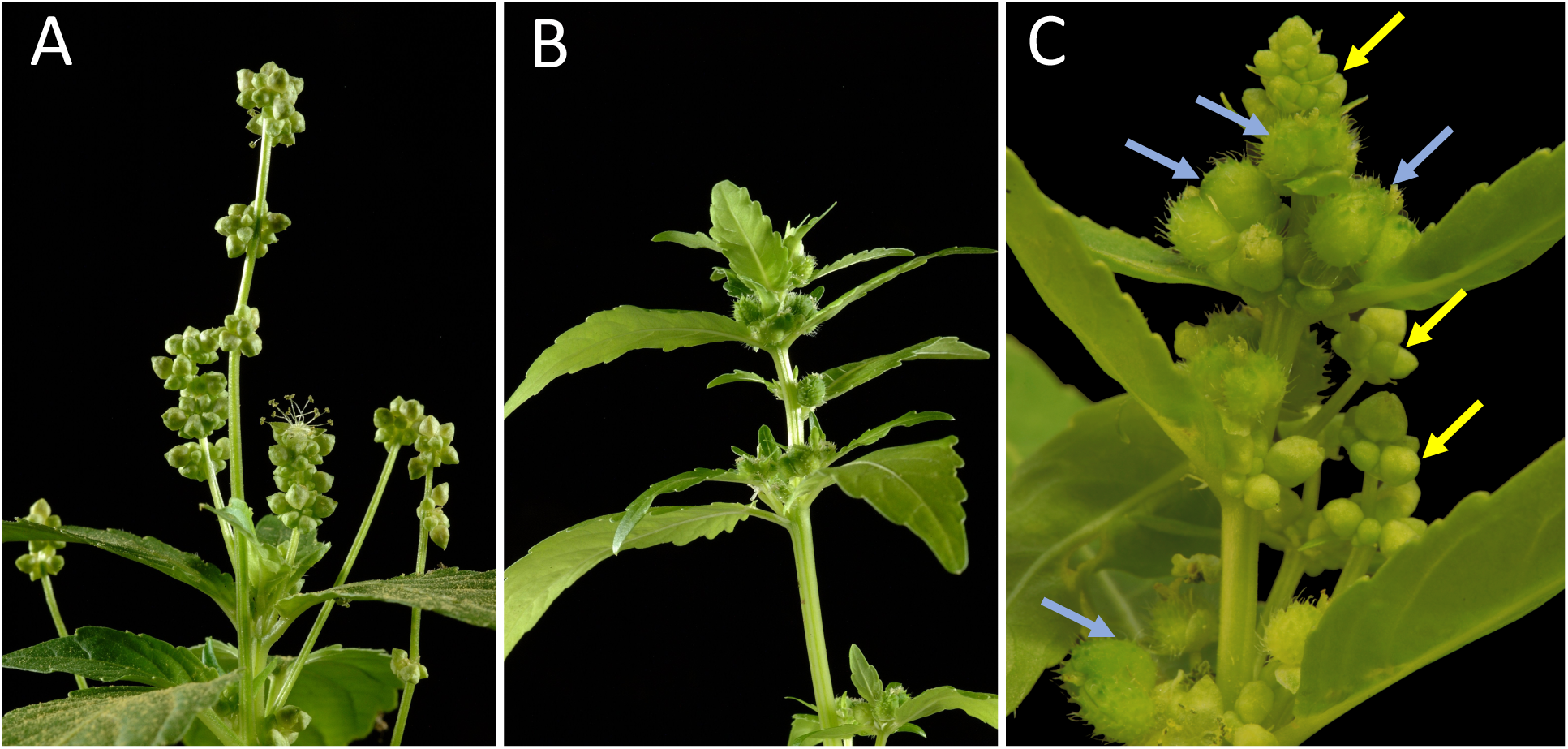
Inflorescence morphology of the different sexes in *Mercurialis annua*. A: Male individual, with erect peduncles bearing male flowers. B: Female individual with female flowers and developing fruits in the leaf axils. C: Female-derived monoecious individual of *M. annua* after four generations of selection on females in the absence of males. Yellow and blue arrows indicate male flowers and female flowers (or fruits), respectively.

We established replicate experimental populations of the diploid lineage of *M. annua* in isolated gardens near Lausanne, Switzerland, and recorded the male flower production of their females growing and evolving in the presence versus absence of males. We established the base population (generation G0) from a single well mixed pool of seed sampled from northeastern Spain. Three ‘control’ populations were established with the typical 1:1 sex ratio of females to males (*15*). In contrast, all males were removed from the other three ‘female-only’ populations. Because males produce large amounts of pollen and mating is effectively random in dense populations of *M. annua* (*16*), all seeds in control populations will have been sired by males, whereas any seeds produced in female-only productions must have been sired by leaky females. We verified that this was so, because the progeny produced in all three control populations had the typical 1:1 sex ratio, whereas those from female-only populations were all female (apart from a very low impact of gene flow from outside the plots, which only occurred in G0; see Methods). We continued the experiment over five consecutive years, from 2012 (generation G0) to 2016 (generation G4). At the end of each spring, after seven weeks of growth and mating, we bulk-harvested seeds from all individuals in each population separately and stored them for the next generation the following spring.

As is common in dioecious plants in general (*17-19*), both females and, more rarely, males of *M. annua* are known occasionally to produce a few flowers of the opposite sex, i.e., they show ‘leaky’ or ‘inconstant’ sex expression (*20*). Accordingly, a fraction of females established as potential parents in the first generation G0 of our experiment also expressed a low degree of leakiness in all populations. Interestingly, females from the base population growing in the field populations in the absence of males, i.e., in the female-only populations of generation G0, produced 2.33 times more male flowers than those growing with males in the control populations (*P* < 0.01; Fig. 2A, C). This difference indicates that leaky sex expression by *M. annua* females has a significant plastic component (*21*). Although plasticity is often interpreted as being adaptive, this has rarely been demonstrated convincingly in the literature (*22, 23*). In our experiment, leaky plasticity was clearly adaptive, because females capable of enhanced leakiness in the absence of males in our experiment enjoyed high fitness by siring all the population’s progeny (Table 1).

**Table 1.** Standardized linear (*β*) and quadratic (*γ*) selection gradients for the male allocation of females grown in the absence of males. Estimates are average linear (*β*) and quadratic (*γ*) selection gradients calculated separately for each experimental population. Standard errors are in parentheses. Stars indicate values that are significantly different from zero at *P* = 0.05, calculated from a comparison of the mean selection-gradient values and the null expectation of zero based on *t*-tests with 2 degrees of freedom (i.e., the experimental populations served as the unit of observation). * : 0.01 *< P <* 0.05 ; ** : 0.001 *< P <* 0.01; *** : *P <* 0.001.

| Generation | Treatment | $\beta$ | $P$-value | $\gamma$ | $P$-value |
| --- | --- | --- | --- | --- | --- |
| **G0** | **Control** | 0.087 (0.048) | 0.21 | - 0.019 (0.084) | 0.19 |
| | **Female-only** | 0.81 (0.043) *** | $2.81 \times 10^{-3}$ | 0.56 (0.052) ** | 0.011 |
| **G1** | **Control** | 0.056 (0.102) | 0.44 | - 0.0006 (0.046) | 0.92 |
| | **Female-only** | 0.73 (0.070) *** | $3.08 \times 10^{-3}$ | 0.26 (0.191) | 0.36 |
| **G2** | **Control** | $9.53 \times 10^{-3}$ ($9.33 \times 10^{-3}$) | 0.22 | - $1.28 \times 10^{-3}$ (0.016) | 0.55 |
| | **Female-only** | 0.89 (0.038) *** | $6.10 \times 10^{-4}$ | 0.44 (0.039) ** | 0.010 |
| **G3** | **Control** | $1.33 \times 10^{-3}$ (0.049) | 0.97 | - 0.104 (0.095) | 0.40 |
| | **Female-only** | 0.72 (0.109) *** | $7.49 \times 10^{-3}$ | 0.30 (0.062) | 0.052 |

**Fig. 2.**
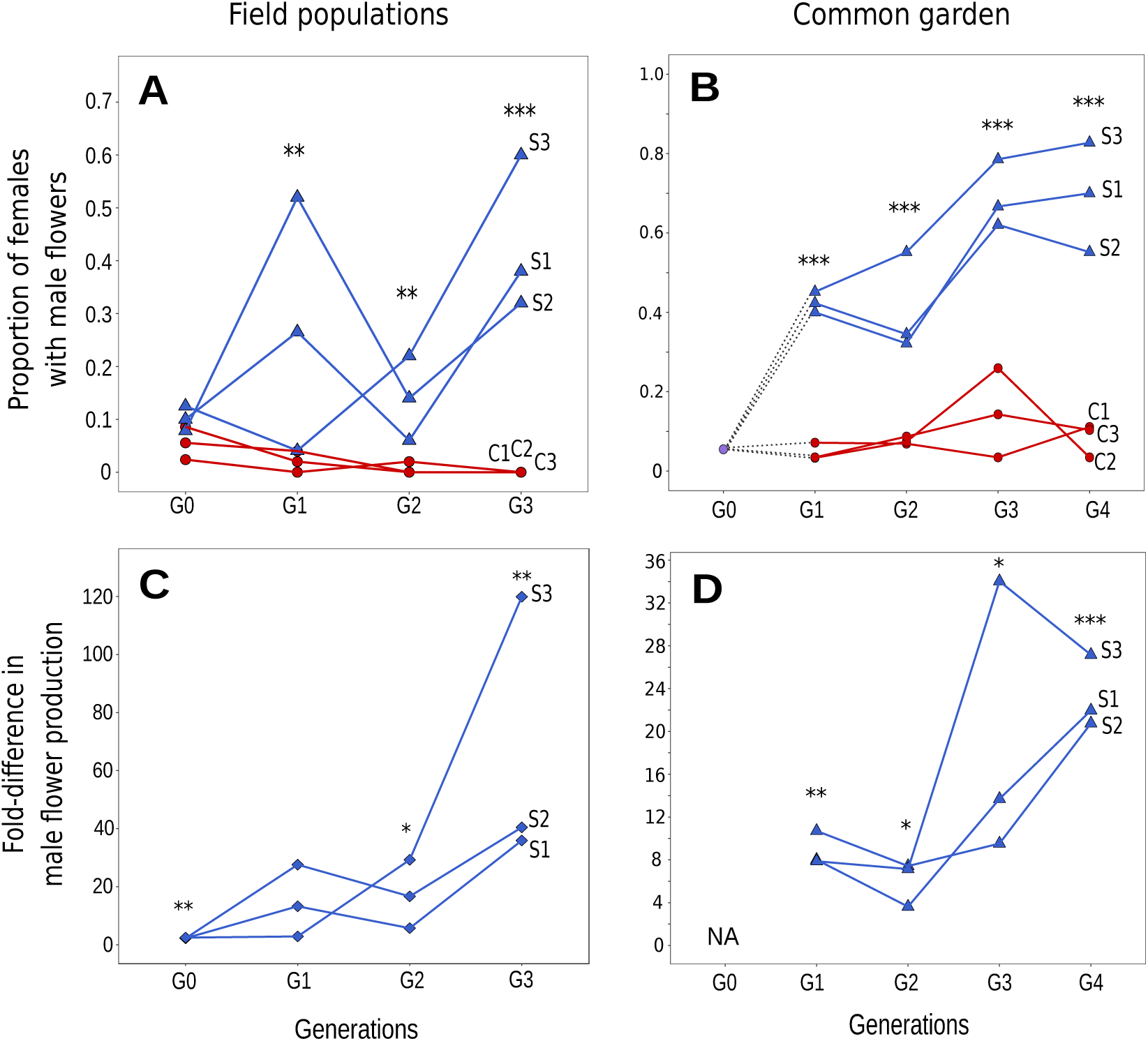
Difference in male-flower production between females in female-only populations and control populations for plants measured in the field populations (A and C) and in the common garden (B and D). A and B: Proportion of females producing male flowers in female-only (blue triangles) and control populations (red circles); i.e., females producing at least as many male flowers as the 95th percentile of male-flower production measured at generation G0, averaged across the three control populations in the field (A), or averaged across all females from the base population in the common garden (B). Stars indicate a significant difference in male allocation between control and female-only populations, tested using a mixed generalized linear model with a binomial distribution, using replicates as random factor. C and D: Difference in male-flower production by females between female-only and control populations, measured as the mean log_2_ fold-difference between the two categories. Stars indicate a log_2_(fold-difference) significantly different from 0, based on a *t*-tests. * : 0.01 *< P <* 0.05; ** : 0.001 < P < 0.01; *** : *P <* 0.001.

We observed a dramatic increase over the course of four generations in both the proportion of females producing male flowers in the female-only populations (Fig. 2A), and in the number of male flowers they produced (Fig. 2C), whereas these changes were not observed in the control populations. Specifically, the proportion of females in the female-only populations that produced more male flowers than the 95th percentile observed in the base population (G0) increased from 5% prior to selection to 69.3% over three generations of selection. On average, females in the female-only populations produced a mean of 14.7 times more male flowers after one generation of selection (i.e., in G1) than females in the corresponding control populations, and a mean of 65.4 times more male flowers after three generations (i.e., in generation G3; Fig. 2C, Table S1). Nevertheless, despite the striking directional changes observed for female-only populations, there were also substantial differences in male-flower production by females among replicate populations in the field, as well as fluctuations among years (Fig. 2A and C). This variation indicates that male-flower production continued to be plastic and potentially sensitive to the environmental conditions experienced among sites. To compare male-flower production in a common environment between females evolving with versus without males, and thus to focus on the evolved response independent of any direct effect of the presence or absence of males, or of other environmental differences between populations, we grew females from each population and each generation (from stored seed) in a large common garden, establishing them in a random block design intermingled with males. Here, too, we observed a substantial and significant increase both in the frequency of females producing male flowers (from 5.48% in the base population G0 to a mean of 69.3% by generation G4; Fig. 2B), as well as in the mean male-flower production of females overall (females in female-only populations produced a mean of 8.77 times more male flowers than those in the control populations in generation G1, a difference that increased to a mean of 23.3-fold by generation G4; Fig. 2D). The observed response by females to selection over four generations corresponds to an increase in male-flower production per biomass of 5.84 standard deviations relative to the base population (Fig. 3). The associated evolutionary rates for female-only populations per generation in our study, calculated in haldanes (*24*), ranged from 0.001 to 0.973, with the highest values recorded in the female-only populations between generations G0 and G1 when the fewest females produced male flowers and selection was accordingly particularly strong (Table S2). Previously observed adaptive responses by plants to natural or artificial selection range from 0 to 0.808 haldanes (*25*). Similarly, responses measured for animals range from 0.001 to 0.74 haldanes (*26, 27*). The evolutionary rates measured in the first generation of our experiment are therefore larger than any reported in the literature, although the rates in subsequent generations lie within the range reported in other studies (*25-27*).

**Fig. 3.**
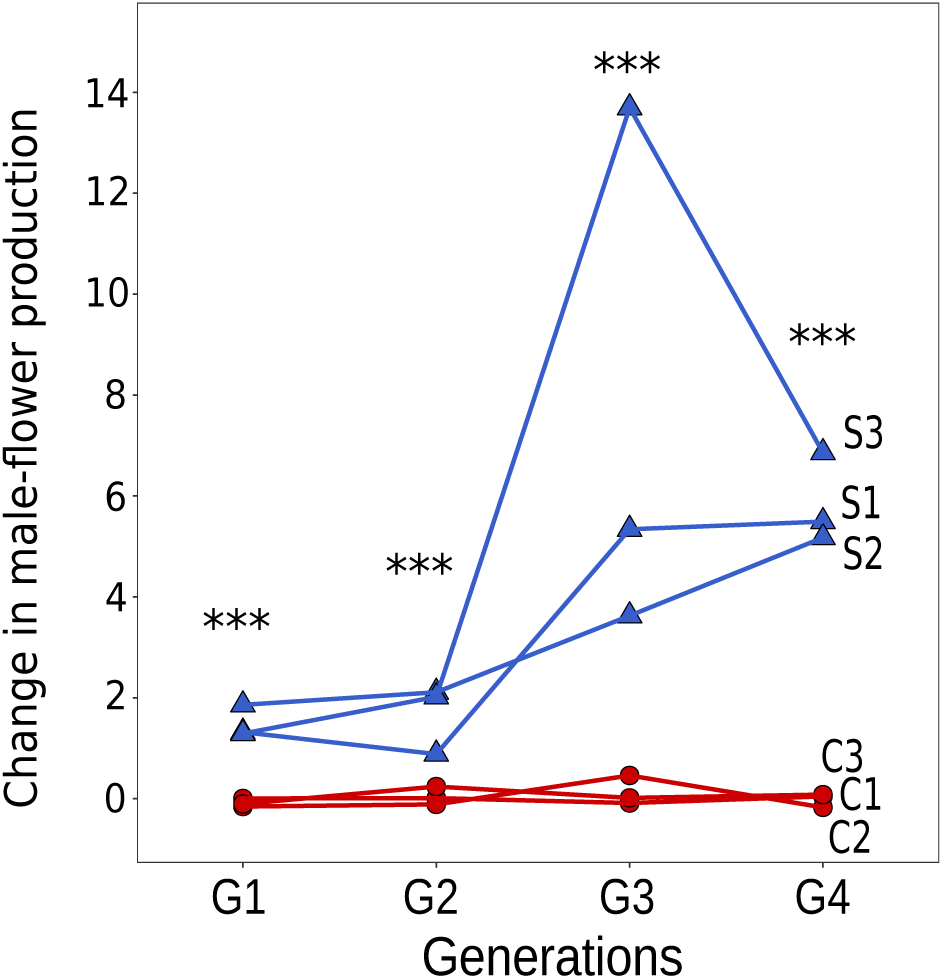
Change over time in male-flower production by females, measured in the common garden in terms of the number of standard deviations (SD) in male-flower production recorded for the base population (also grown in the common garden). Stars indicate a significant difference in mean male-flower production between selected and control lines, based on permutation *t*-tests with 100,000 bootstraps and two degrees of freedom: *** : *P <* 0.001.

**Fig. 3.**
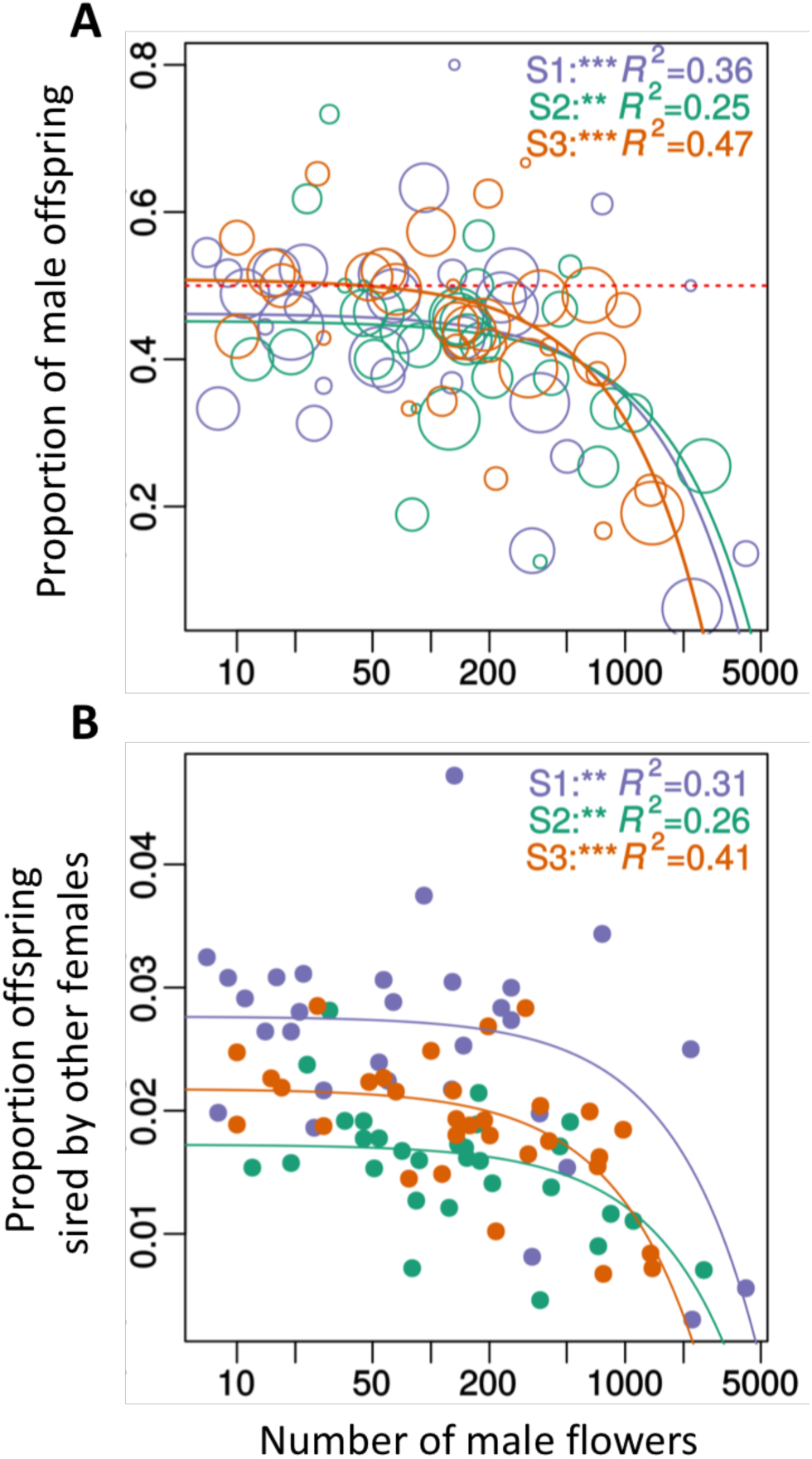
Maternal progeny sex ratios and estimation of the proportion of outcrossed seeds of females from experimental populations in the presence of males in low-density arrays. a, Proportion of male progeny as a function of the number of male flowers produced by maternal females. Linear models were weighted by binomial confidence intervals inferred from the number of male and female progeny in each family. Each circle represents one maternal family, the size being proportionate to the reciprocal of binomial confidence intervals. b, Estimated proportion of offspring sired by other females. Each point represents one maternal family. Colours correspond to the experimental population replicates of origin of the maternal plants (Purple: S1, Green: S2, Orange: S3). Note that regression lines are actually linear and the curvature is caused by the log-scaled X-axis. Stars in the upper of panels indicate significance of regression estimates for each of the experimental populations *: 0.01 *< P <* 0.05; **: 0.001 < *P* < 0.01; ***: *P <* 0.001, R^2^ indicates the proportion of variance explained by the linear models.

High estimates of evolutionary rates, like those found in our study, typically characterize contemporary evolution at ecological time scales, which is expected to be ‘rapid’ in comparison to rates over longer period of time that may experience reversal and stasis rather than only strong directional selection (*28, 29*). The rapid response to selection in our experiment not only reflects intense competition for siring success among females deprived of males and a likely close correspondence between mating and male-flower production in *M. annua* (*30*), but also high heritability of the selected trait. We estimated the realized heritability of male-flower production by females through inversion of the breeders’ equation (*31*) as *h*^2^ = 0.73 (SD = 0.48). This value is similar to that typical of more canalized traits such as floral diameter (e.g., (*32-34*)), but is much higher than that typically recorded for less canalized traits such as flower number (e.g., *h*^2^ = 0.04 in *Silene latifolia*; (*34*)). Importantly, the high heritability of leaky sex expression observed in *M. annua* indicates that leakiness in sex expression in dioecious plants (*17*) cannot generally be attributed to developmental instability and may often be both responsive to selection and a fundamental capacitor for the breakdown of dioecy (*9, 35*).

The genetic architecture of variation in male-flower production by females in *M. annua* is not yet known, but the loci responsible are clearly not found exclusively on the Y chromosome, which is missing in the female-only populations. However, females of *M. annua* with an evolved male function produce their male flowers in sub-sessile axillary inflorescences that are typical of natural females (compare Figs. 1B and 1C), rather than on erect inflorescence stalks that are characteristic of XY males (Fig. 1A) and that promote better pollen dispersal (*30*). Thus, whereas a Y-linked promotor of male function directly governs male versus female inflorescence differentiation in *M. annua*, male-flower production is governed by loci that are not themselves Y-linked. It follows that selection on standing genetic variation in our experiment has evidently favored the masculinization of gene expression of loci downstream of a trans-acting sex-determination switch. Experimental phenotypic masculinization of female *M. annua* has been achieved via exogenous application of plant growth hormones, which likely play a key role in sex expression in *M. annua* (*36*), as in dioecious plants more generally (*37*).

We next determined the extent to which the increased production of male flowers by females in female-only populations has altered the mating system of their populations. Using polymorphism at eight microsatellite loci (*38*), we estimated the rate of self-fertilization, *s*, in all six experimental populations at generations G1 and G4. Whereas all dioecious control populations were completely outcrossing at both generations G1 and G4, *s* was significantly greater than zero in all three female-only populations at generation G1 (mean *s* = 0.19; Table 2) and in two of them at G4 (mean *s* = 0.12). Thus, a significant proportion of the progeny produced in female-only populations were self-fertilized, confirming the ability of monoecious individuals of *M. annua* to benefit from reproductive assurance in the absence of mates (*39*). Nevertheless, in the female-only populations, most progeny were the result of outcrossing, confirming that there is strong competition for outcross siring success among females with an evolved male function under the dense growing conditions of our experiment.

**Table 2.** Selfing rates inferred for the experimental populations. Selfing rates (s) were calculated at generations G1 and G4 on the basis of eight polymorphic microsatellite markers using RMES(*43*). 95% confidence intervals are indicated. *P*-values correspond to test for whether s was significantly different from 0 (ns: non-significant at *P* < 0.05).

| Generation | Treatment | Population | Sampling size | *s* | *CI*<sub>95</sub> | *P*-value |
| --- | --- | --- | --- | --- | --- | --- |
| G1 | Control | C1 | 114 - 120 | 0.047 | [0, 0.14] | ns |
|  |  | C2 | 111 - 120 | 0.000 | [0, 0.038] | ns |
|  |  | C3 | 103 - 120 | 0.000 | [0, 0.062] | ns |
|  | Female-only | S1 | 116 - 120 | 0.124 | [0.042, 0.21] | < 0.001 |
|  |  | S2 | 117 - 120 | 0.225 | [0.13, 0.31] | < 0.001 |
|  |  | S3 | 118 - 120 | 0.225 | [0.11, 0.33] | < 0.001 |
| G4 | Control | C1 | 116 - 120 | 0.019 | [0.00, 0.078] | ns |
|  |  | C2 | 119 - 120 | 0.000 | [0.00, 0.055] | ns |
|  |  | C3 | 114 - 120 | 0.062 | [0.00, 0.16] | ns |
|  | Female-only | S1 | 118 - 120 | 0.172 | [0.053, 0.28] | < 0.005 |
|  |  | S2 | 120 | 0.074 | [0.00, 0.16] | ns |
|  |  | S3 | 117 - 120 | 0.121 | [0.016, 0.23] | < 0.005 |

Finally, we asked whether females with an enhanced male function would also be capable of siring progeny in competition with normal XY males, which produce much more pollen and have inflorescences that are well adapted to pollen dispersal (*30*). To address this question, we established six separate common gardens (‘mating arrays’) from seeds produced in the experimental populations after a further generation of selection (i.e., G5), re-introducing males from the control populations into the female-only populations, such that all common gardens comprised a lattice with 50% males. Family sex ratios from the gardens established with seed from control populations did not differ from 50% (mean of 50.3%; binomial test: *P* = 0.551, Table S3), as expected for a dioecious sexual system with obligate outcrossing and males as the only sires. In contrast, families from gardens established with females from the female-only populations produced an average of 43.6% males in their progeny, significantly lower than 50% (binomial test, *P* < 0.001; Fig. 3 and Table S3). Because XY males should sire an equal number of sons and daughters, the 6.4% male deficit in the progeny of females evolved in the female-only populations implies that 12.8% of progeny (6.4% × 2) had been sired by females. The fact that females with more male flowers also produced more daughters (Fig. 4A) indicates that the lower frequency of sons in their progeny was partially a result of self-fertilization (Fig. 3B).

Taken together, our results demonstrate that plants from a species displaying leaky dioecy can rapidly evolve functional hermaphroditism (monoecy) if deprived of their opposite-sex partners. In the case of *M. annua*, the masculinization of females in the absence of males is sufficient to alleviate pollen limitation, to influence the mating system and, importantly, to shift the sexual system away from dioecy to functional monoecy. In the continued absence of males, hermaphroditism should become further stabilized with an increasing allocation of resources to pollen production (*40*). In contrast, the re-introduction of males into such modified populations, which in nature could occur by dispersal from populations that have remained dioecious (and as illustrated by our male reintroduction experiment), should establish functional androdioecy (a sexual system in which males are maintained with hermaphrodites) (*11, 41*). Androdioecy is thought to evolve in response selection when females become separated from males at low density or during colonization (*42*). Our study thus provides not only the first experimental demonstration of a mechanism for the evolution of functional hermaphroditism from dioecy, but also experimental validation of the steps thought to promote the evolution of androdioecy.

## Acknowledgments

N. Geldner, P. Meystre, A. Wolf, A. Guisan and D. Wagener provided garden space, Nicolas Ruch and Aline Revel helped with plant care, and D. Savova-Bianchi, T. Martignier, L. Santos, W-J. Ma, P Veltsos, A-P Machado, F. Gréverath, C. Dupuis, C. Jaulin, I. El M’Ghari, R. El Assad, S. Beuvier, S. Tesson and L. Olazcuaga helped with plant genotyping and phenotyping. We thank Tad Kawecki, Gabriel Marais and Brian Hollis for valuable discussion.

## Author contributions

Designed the study: JRP and GGC. Performed the experimental work: GGC, XL, JFG, and JRP. Performed the mating-system analysis: YC. Wrote the manuscript: JRP, GGC, XL, JFG. All authors approved the final version of the manuscript.

## Data and materials availability

All data will be uploaded to the Dryad repository upon final publication.

## Materials and Methods

### Experimental design of the selection experiment

Our experiment compared the evolution of female phenotypes growing in the presence versus absence of males. We established six experimental populations of dioecious *Mercurialis annua* in isolated locations on the campus of the University of Lausanne, and in private gardens distributed around the city. Three ‘control’ populations had the natural 1:1 sex ratio usually found in the field (and in our starting seed population), while three ‘female-only’ populations were established without males. The populations were derived from an experimental seed bank (base population G0) established by pooling seeds of open pollinated plants from 25 demes of a metapopulation in northern Spain (*20*).

To establish the experimental populations each year, we raised seedlings from seeds in seed trays in a mixture of horticultural soil and perlite, and let them grow in a glasshouse until their sex could be determined at the onset of flowering. In the first generation (G1, spring, 2012), 180 plants were allocated in a random fashion to each of the three control and female-only populations. In subsequent generations, population sizes were increased to 210 plants per population. The site allocated to each replicate was randomized for each of the first three generations. From generation four, we maintained experimental populations at the same locations each year to avoid potential gene flow from control to treatment plots via plants that might have established in the habitat from previous generations. We nevertheless surveyed sites carefully each year and removed any stray individuals. Sex in *M. annua* is genetically determined by an XY sex-chromosome system^18,19^. Female-only populations in our experiment had no Y chromosomes, so that males in their progeny could only be attributable to pollen migration from fathers outside the population. We found very few such males (a mean of 1.33%) in the first-generation (G1) progeny of the three female-only populations, which we removed. No genetic (XY) males were observed in subsequent generations. Plants were established at the experimental sites in 3.5 litre pots with soil substrate Ricoter 140® and three individuals per pot (in the control population, either two males and a female, or two females and a male, in alternating fashion). Pots were watered once daily with an automatic watering system. Individuals were allowed to mate freely in their experimental sites for approximately seven weeks each spring. They were harvested and measured. Seeds for each subsequent generation were sub-sampled from the bulk of the previous year’s production for that population, and all remaining seeds were stored in air-tight containers at 4° C for later use in the common garden.

### Harvest protocol and phenotype measurements

We recorded the phenotypes for the plants growing in the experimental populations each generation, as well as in a common garden after four generations of selection (i.e., in early summer, 2016). At the time of harvest each year, we recorded the following traits on 50 randomly chosen females per replicate: plant height; above-ground biomass; number of male flowers on the plant (before anther dehiscence); and number and weight of seeds produced. For the common garden, plants from each generation and replicate were established in a random block design at the same high density experienced during the experiment. The common garden thus comprised 24 population × generation combinations, with 30 females from each population and generation, 20 males from each control population and generation (generations 1 through 4), and 90 females and 60 males from the initial seed pool (G0). The design comprised 10 blocks, with a total of 30 males and 81 females distributed evenly across blocks, with all population × generation combinations represented in each of 10 blocks. Because the plant density in the garden was kept high, seed production by females was not pollen-limited (*39*). We measured the size of all plants and components of their reproductive allocation after seven weeks of growth.

For each individual of the common garden, we recorded the same traits as in the field replicates, as well as weight of male flowers produced by males. For all plants measured, both from field replicates and common-garden data, we computed male allocation as the ratio of male flower biomass over the above-ground biomass. To compute the total biomass of male flowers when it was too low to be measured by our balance (which was precise to 0.00001 g), we calibrated the average mass of a single male flower from 35 females from the common garden. Similarly, we computed female allocation as the total seed biomass over above-ground biomass, as well as the ratio of height to biomass. We also computed the mean degree of sex ‘inconstancy’ of females, *d* (i.e., the proportional extent to which females’ male flower production approached that of pure males), as the mean male allocation for females divided by the mean male allocation for males growing under similar conditions (*20*). Means were calculated across females of each population separately, relatively to males in control populations from the same generation.

### Statistical analysis of selection gradients and heritability

We calculated the prospective relative fitness for females in each of the replicate populations in the field following Morgan & Schoen (*44*) and Dorken and Pannell (*40*). Specifically, we estimated directional (*β*) and nonlinear selection gradients (*γ*) as the linear and quadratic regression coefficients of male allocation on relative fitness (*45*). Quadratic selection gradients were calculated as twice the quadratic regression coefficients, as recommended by Stinchcombe *et al*. (*46*). We used Mann-Whitney ranked tests to compare male allocation, female allocation and above-ground biomass of females between female-only and control populations, at each generation, for both field replicates and the common garden. The phenotypic response to selection was calculated from females in the common garden as the mean difference in male allocation between generation four and the base population (*Δ*_*a*_), measured in haldanes (*Δ*_*a*_/*V*), where *V* is the pooled within sample variance in consecutive generations (*24*), as well as the number of standard deviations for male allocation in the initial generation over which the mean male allocation was shifted by selection (*Δ*_SD_) (*24*).

We estimated the selection differential, *S*, as the covariance between male allocation and relative fitness, based on plants measured for each population in the experimental gardens. Using the breeder’s equation (*31*), we calculated the narrow-sense realised heritability as *h*^*2*^ = *Δ*_*a*_*/S*, averaged across the three female-only populations after the three last rounds of selection. Values for the response to selection (*Δ* _*a*_) used in the calculation of *h*^*2*^ were taken from populations in the field (to match the selection differential estimates). We averaged across the three years to estimate *h*^*2*^ for male allocation.

### Mating-system estimation

To assess the extent to which the evolution of a male function in females led to a change in their mating system from outcrossing to partial selfing, we used microsatellites to estimate the selfing rate of females in all populations at generations G1 and G4, and in the original seed pool used (generation G0). Sample sizes ranged from 103 to 120 progeny per experimental population and generation, genotyped at eight microsatellite loci (*38*). In addition we also genotyped 55 to 62 individuals from the initial G0 seed pool, depending on microsatellite locus. Seeds for genotyping were sown in trays with a mixture of horticultural soil and perlite, and plants were raised to maturity over about five weeks. Leaf material was collected from 120 individuals per population and generation and dried in a drying cabinet at 55 degrees for 48h. After homogenization using metal beads and TissueLyser, individual total DNA was extracted from leaf material from each individual using BioPrint 96 (Qiagen, Germany), following the manufacturer’s recommendations. PCR amplifications were performed in 10 µL reactions, using GoTaq DNA polymerase (Promega), following the protocol in Machado et al. (*38*). Fragment lengths were analysed with internal size standard GeneScan-350 LIZ using ABI3100 Genetic Analyzer (Applied Biosystems), and were scored with GeneMapper v4.0 (Applied Biosystems).

We used the software RMES to estimate selfing rates for each replicate, using the maximum likelihood unconstrained method (*43*). RMES computes selfing rates (*s*) from the multilocus structure of microsatellite variation, taking into account the possible contribution of null alleles to estimates of homozygosity. We also used RMES to test whether the estimated selfing rates differed significantly from zero (ML constrained *s* = 0; *P <* 0.05).

### Assessment of siring success and the sexual system

We established female progeny from the fourth generation of selection from each population, i.e., the fifth generation of our selection experiment (collected in 2016) and used the sex ratios of their progeny to determine their ability to sire progeny in competition with males, which we reintroduced to the female-only populations from the control populations at a frequency of 50%. Seeds were first germinated in seed trays and raised to maturity under greenhouse conditions at the University of Lausanne before being transplanted into 14 cm-diameter pots with substrate Ricotor 140® and 5g/L slow-release fertiliser (Tardit 6M). We then set up six mating arrays at the nursery site of the Ville de Lausanne, Bourdonnette, one array for each of the six experimental populations. In each array, we arranged 50 females and 50 males in a 10 × 10 lattice at 1 m intervals. Females in a given array were sampled from one of the six respective experimental populations (three control and three female-only populations), while males were sampled from a single pool of males sampled in equal proportions from each of the three control populations.

Plants were harvested after two months of open pollination. For each female, we measured its height, the number of male flowers, the dry biomass and the number of seeds produced. We also measured the biomass and the biomass of male flowers produced by each of a random sample of 10 males from each array. Male flower number was estimated by calibrating against drymass of male flowers. We estimated the sex ratio in the progeny of each female by raising up to 200 progeny from seed (depending on numbers available on the plant) in seedling trays with soil substrate Ricoter 153®. We recorded the sex of each seedling as it began flowering, over a period of approximately eight weeks.

### Model for estimating the paternity of seeds sired in mating arrays

Within a mating array, ovules on a focal female may receive pollen from three different sources: (1) self-pollen from the focal plant itself (if it produces pollen); (2) outcross pollen from surrounding male plants; and (3) outcross pollen from surrounding evolved females. We assumed a mass-action model, such that the fate of each ovule is determined by probabilities associated with these three pollen sources. Under this model, the proportions of self-sired *s*, male-sired *m* and female-sired seeds *f* are given, respectively, by:

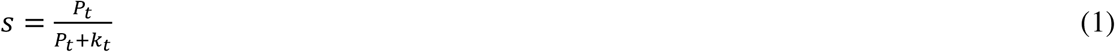

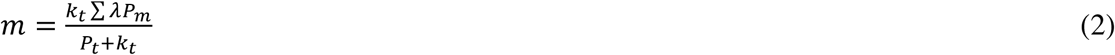

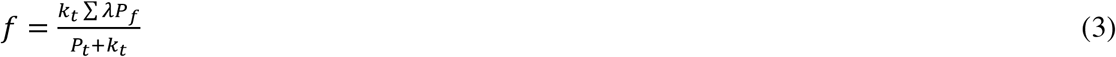

where *P*_*t*_, Σ*P*_*f*_ and Σ*P*_*m*_ are the number of male flowers produced by the target female, the surrounding females, and the surrounding males in the array, respectively, *k*_*t*_ is a dispersion factor calculated for each focal plant that reduces the contribution of outcrossed pollen as a function of its distance from the focal plant, *λ* is a constant that accounts for the better pollen dispersal of males relative to females as a result of their erect inflorescence morphology (estimated to be 1.6; (*16, 47*)). For males, we transformed male flower dry mass into flower counts based on the coefficient 0.0002 g per male flower. Note that the location of the focal plant in the array had no significant effect on variation in the proportion of males in the progeny, possibly due to the great difference in the amount of pollen produced by males and females, so we modelled mating in terms of pollen clouds that took no account of position.

Because *M. annua* is a species with an XY sex-determining system, only seeds sired by males will result in male progeny. Thus, *m* in equation (2) will be twice the proportion of male plants in each family. We solved equation (2) for *k*_*t*_ and used it to estimate the proportion of self- and female-sired seeds for each family.

---

**Table S1.** Means and standard deviations of phenotypic traits of females in the common garden, averaged across replicates per generation. N: sample size; V: height/drymass; Male allocation: mass of male flowers/plant drymass. Female allocation: mass of seeds/drymass. ***d***: degree of inconstancy. For all statistics, standard errors are shown in parentheses. Student’s *t-*tests (two-sided) were used for statistical comparisons of females between control and female-only populations, with experimental populations as the unit of replication. * : 0.01 *< P <* 0.05 ; ** : 0.001 *< P <* 0.01 ; *** : *P <* 0.001.

| Generation | G0 | G1 |  |  | G2 |  |  |
| --- | --- | --- | --- | --- | --- | --- | --- |
|  | 50%<br>males | 50%<br>males | Males<br>removed | *P* | 50%<br>males | Males<br>removed | *P* |
| N | 73 | 87 | 87 | - | 79 | 86 | - |
| Drymass (g) | 8.80 (4.41) | 9.80<br>(2.95) | 11.69<br>(3.49) | 0.42 | 9.68<br>(2.97) | 10.47<br>(2.95) | 0.24 |
| V (cm/g) | 8.43 (4.14) | 7.34<br>(2.16) | 5.98<br>(1.90) | 0.39 | 6.93<br>(1.91) | 6.39<br>(1.88) | 0.14 |
| Male allocation<br>(x10<sup>-4</sup>) | 1.58 (5.44) | 1.10<br>(2.69) | 9.66<br>(14.1) | 9.9<br>x10<sup>-3</sup> ** | 1.81<br>(4.10) | 10.6<br>(16.7) | 0.046 * |
| Female<br>allocation | 0.11 (0.031) | 0.11<br>(0.030) | 0.13<br>( 0.036) | 8.15<br>x10<sup>-3</sup> *** | 0.12<br>(0.036) | 0.13<br>( 0.035) | 0.31 |
| *d* (x10<sup>-3</sup>) | 0.90 (3.08) | 0.68<br>(1.73) | 5.92<br>(8.76) | 2.55 x10<sup>-13</sup><br>*** | 1.02<br>( 26.5) | 6.21<br>(11.0) | 2.03 x10<sup>-13</sup><br>*** |
| Generation | G3 |  |  | G4 |  |  |  |
|  | 50%<br>males | Males<br>removed | *P* | 50% males | Males<br>removed | *P* |  |
| N | 84 | 87 | - | 76 | 88 | - |  |
| Drymass (g) | 10.74<br>(3.67) | 10.95 (3.30) | 0.79 | 8.99 (3.14) | 10.79<br>(3.86) | 0.36 |  |
| V (cm/g) | 6.48<br>(2.07) | 6.19 (1.70) | 0.61 | 7.33 (2.15) | 6.86 (2.71) | 0.66 |  |
| Male allocation<br>(x10<sup>-4</sup>) | 2.27<br>( 4.04) | 42.6 (57.3) | 0.14 | 1.48 ( 3.08) | 33.3 (45.4) | 6.68 x10<sup>-3</sup><br>** |  |
| Female<br>allocation | 0.12<br>(0.034) | 0.11 ( 0.041) | 0.61 | 0.13 (0.033) | 0.12<br>( 0.042) | 0.022 * |  |
| *d* (x10<sup>-3</sup>) | 13.4<br>(27.9) | 25.2 (41.4) | < 2.2 x10<sup>-16</sup><br>*** | 0.83 (1.94) | 19.2 (261) | < 2.2 x10<sup>-16</sup><br>*** |  |

**Table S2.** Evolutionary rates in haldanes, measured per generation, separately for each of the replicate control (C1-C3) and female-only (S1-S3) populations. NA: non-available data for the last generation measured in the field (no generation 5 in the field has been recorded).

| Populatio<br>n | Treatmen<br>t | G1 |  | G2 |  | G3 |  | G4 |  |
| --- | --- | --- | --- | --- | --- | --- | --- | --- | --- |
|  |  | Garde<br>n | Field | Garde<br>n | Field | Garde<br>n | Field | Garde<br>n | Fiel<br>d |
| **C1** | Control | 0.001 | $\bar{-}$<br>0.459 | 0.011 | 0.055 | -0.220 | 0.056 | 0.267 | NA |
| **C2** | Control | -0.186 | $\bar{-}$<br>0.337 | 0.126 | $\bar{-}$<br>0.079 | 0.593 | $\bar{-}$<br>0.200 | -0.682 | NA |
| **C3** | Control | -0.122 | $\bar{-}$<br>0.215 | 0.343 | $\bar{-}$<br>0.326 | -0.222 | 0.057 | 0.093 | NA |
| **S1** | Female-only | 0.973 | 0.536 | -0.257 | $\bar{-}$<br>0.715 | 0.894 | 0.671 | 0.022 | NA |
| **S2** | Female-only | 0.951 | 0.601 | 0.056 | $\bar{-}$<br>0.513 | 0.285 | 0.262 | 0.214 | NA |
| **S3** | Female-only | 0.841 | $\bar{-}$<br>0.297 | 0.286 | 0.548 | 0.853 | 0.857 | -0.453 | NA |

**Table S3.** Number of female and male parents and the total number of male and female progeny assayed for each mating array. Results are presented separately for each of the replicate female-only (S1-S3) and control populations (C1-C3) and for all female-only and control populations combined. Binomial tests were performed for the observed progeny sex ratios against the null hypothesis of an unbiased sex ratio (0.5).

|  | No of females | No of males | Male progeny | Female progeny | Progeny sex ratio | Binomial test |  |
| --- | --- | --- | --- | --- | --- | --- | --- |
|  |  |  |  |  |  | P value | 95% CI |
| **S1** | 33 | 35 | 873 | 1130 | 0.4358 | < 0.001 | 0.414 - 0.458 |
| **S2** | 35 | 41 | 838 | 1165 | 0.4184 | < 0.001 | 0.397 - 0.440 |
| **S3** | 33 | 42 | 938 | 1129 | 0.4548 | < 0.001 | 0.432 - 0.476 |
| **All female-only populations** | 101 | 118 | 2649 | 3424 | 0.4362 | < 0.001 | 0.424 - 0.449 |
| **C1** | 46 | 48 | 1820 | 1762 | 0.5081 | 0.341 | 0.492 - 0.525 |
| **C2** | 32 | 44 | 1510 | 1564 | 0.4912 | 0.339 | 0.473 - 0.509 |
| **C3** | 33 | 43 | 1426 | 1371 | 0.5098 | 0.307 | 0.491 - 0.529 |
| **All control populations** | 111 | 135 | 4756 | 4697 | 0.5031 | 0.551 | 0.493 - 0.513 |

